# Coordination between endoderm progression and gastruloid elongation controls endodermal morphotype choice

**DOI:** 10.1101/2023.02.07.527329

**Authors:** Naama Farag, Chen Schiff, Iftach Nachman

## Abstract

Embryos mostly follow a single morphogenetic trajectory, where variability is largely quantitative with no qualitative differences. This robustness stands in contrast to in-vitro embryo-like models, which, like most organoids, display a high degree of variability. What makes embryonic morphogenesis so robust is unclear.

We use the gastruloid model to study the morphogenetic progression of definitive endoderm (DE) and its divergence. We first catalog the different morphologies and characterize their statistics. We then learn predictive models for the lineage morphotype based on earlier expression and morphology measurements. Finally, we analyze these models to identify key drivers of morphotype variability, and devise personalized (gastruloid-specific) as well as global interventions that will lower this variability and steer morphotype choice. In the process we identify two types of coordination that are lacking in the in-vitro model but are required for robust gut tube formation.

We expect the insights obtained here will improve the quality and usability of 3D embryo-like models, chart a methodology extendable to other organoids for controlling variability, and will also shed light on the factors that provide the embryo its morphogenetic robustness.

## Introduction

Embryos of a given species and sex generally follow a single morphogenetic trajectory, from zygote to the end of organogenesis. Variation is quantitative (organ length, number of cells) within a certain range, but not qualitative. For example, all mouse embryos develop a single gut tube, stretching along their antero-posterior axis, and a fixed number of bilaterally-arranged somites arranged symmetrically along this axis (1). Moreover, even in the face of external perturbations (mechanical compression, cell ablation or addition), embryos demonstrate morphogenetic robustness (2,3). The processes driving the morphogenetic dynamics leading to such patterns have been extensively studied in embryos, yet it is not clear what makes them so robust (1,4–10).

In recent years, in-vitro multi-cellular embryo-like models such as embryoid bodies and gastruloids have been used to dissect many aspects of early embryonic development (11–14). Given proper conditions, gastruloids will differentiate in a partially organized manner, break their radial symmetry and develop an antero-posterior (AP) axis (11,15,16). Under specific signal conditions, they can also display trunk elongation (11,13), and even later structures such as somites, gut tube and neural tube (17,18). These in-vitro models, in contrast to embryos, are highly variable, both in terms of the efficiency of forming specific structures, the timing of their formation, and the resulting morphologies obtained (11,19–21). For example, gastruloids can form endodermal gut-tube like structures at varied cell-line dependent efficiencies, but can generate other endodermal morphologies of different sizes, orientations and shapes (13,18,22,23). This degree of phenotypic variability, had it existed in embryos, would have resulted in high rates of lethality or congenital malformations. It is currently not clear what gives the embryo its morphogenetic robustness, nor what is lacking in in-vitro models to obtain such robustness at equivalent stages. For the in-vitro models, this poses a key limitation to fully exploiting their scientific and translational potential. It is therefore pivotal to quantify the extent of phenotypic variation, to identify its drivers, and to leverage this knowledge for data-driven strategies to control and curb morphogenetic variability.

Here we use gastruloids and the progression of definitive endoderm (DE) in them to study divergence of morphogenetic programs, employing a dual Bra-GFP and Sox17-RFP reporter cell line (24). We characterize and quantify the possible endoderm morphologies in gastruloids. We then use a machine learning approach to predict endoderm morphology from early gastruloid shape and mesoderm/endoderm progression. Finally, we analyze the learned predictors to identify the key affecting parameters, and devise interventions that divert the morphology distribution towards the desired outcomes.

## Results

### Gastruloid Endoderm displays distinct classes of morpho-types

To study the drivers of endoderm morphology in gastruloids, we employed a dual reporter mESC line, Sox17-RFP/Brachyury-GFP, in order to track in real time the onset, expansion and morphology of mesoderm and endoderm in developing gastruloids (24).

We first set out to characterize the possible DE domain morphologies that are obtained during gastruloid development. Gastruloids were differentiated using the standard protocol, which includes aggregation in N2B27 media for 48 hrs, a CHIR pulse between 48-72h post-aggregation, then back to N2B27 media for the rest of differentiation (Fig. 1a) (also in Methods). In some experiments we extended the differentiation protocol up to 120hrs using the trunk-like-structure protocol (5% Matrigel between 96hrs-120hrs (18)). The gastruloids were live imaged during the differentiation process, starting from 48 hrs up until 96 or 120 hrs.

**Figure 1.**
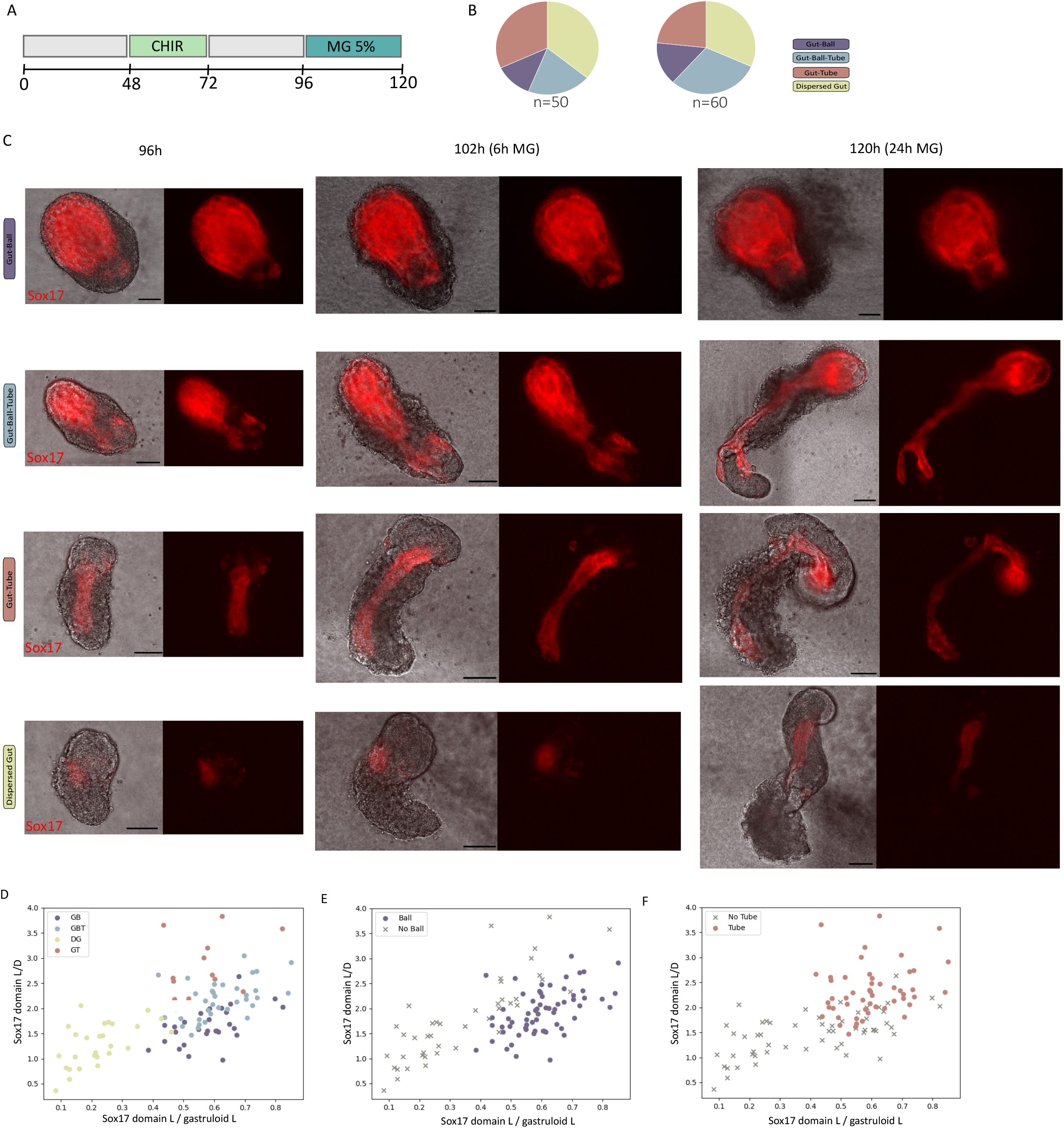
Gastruloid endoderm displays distinct classes of morpho-types. **A**. Differentiation time-line. **B**. Distribution of DE morphologies, at 96h, in two experimental repeats. **C**. Sample images of gastruloids with different DE morphotypes at 96, 102 and 120hrs. **D-F**. Representation of the gastruloids according to dimensionless morphological parameters measured at 96h. Gastruloid data points are colored by the four morpho-types (D), or by binary appearance of Ball (E) or Tube (F) in their morphology.

Using this protocol, we observed that endodermal cells within the gastruloid display different morphologies along the protocol timeline. At 96 hrs these morphologies can be divided into four major *morphotypes*: ‘Gut-Ball’ (GB), ‘Gut-tube’ (GT), ‘Gut-ball-tube’ (GBT) and ‘Dispersed Gut’ (DG) (Fig. 1b,c). After gastruloid elongation, the identity of each morphotype becomes more distinct (Fig. 1c). The *GB* morphology appears as a ‘Ball’ of tangled lumens composed of Sox17(+) cells, which remains at one pole of the gastruloid upon its elongation (22) (Movie S1).

*GT* morphology appears as a narrow tube of Sox17+ cells that elongates with the gastruloid along the A-P axis (13,18) (Movie S2). The GT and GB morphologies are not mutually exclusive. Some gastruloids contain a GB structure, with a GT elongating from it along the A-P axis (Movie S3). This morphology is defined as ‘Gut-Ball-Tube’ (or *GBT*). The fourth morphology (DG) encompasses gastruloids that lack a defined morphology of Sox17+ cell groups.

The different morphologies can be continuously quantified within a phase-space of simple morphometric parameters at the 96 hrs time point, such as endoderm domain length/diameter, or ratio of endoderm domain length to gastruloid length (Fig. 1d). This separation can also be seen when looking on binary appearances of Ball or Tube (Fig. 1e,f). As can be seen, the morphometric parameters do not perfectly distinguish between the 4 morphotypes (or even the binary appearance of ball or tube), but do provide an objective approximate description for them.

### Early gastruloid morphology predicts endoderm morpho-type

The results show that different morphotypes arise in a single experiment, where all gastruloids receive the same treatment (Fig. 1a). This gastruloid-to-gastruloid variation can stem from multiple factors, such as seeding, aggregation, cell viability and cell state differences between the gastruloids. We therefore set out to analyze the dependency of morphotype choice on early measurable parameters along the differentiation process. The rationale is that if enough relevant parameters are measured, morphotype choice may be predicted in each such experiment, and the predictive parameters may later help to control or enrich for specific morphotypes.

In order to understand the gastruloid internal parameters affecting DE morphotype choice, we have collected morphology and timing parameters from each gastruloid, both at the specific lineage level (mesoderm and endoderm) as well as the whole gastruloid level. These parameters were evaluated at several time points between 48 and 72 hours (Figure S1). Some of these parameters (such as gastruloid initial size) can be externally controlled to some degree of accuracy, while others (such as gastruloid morphology parameters) are not directly controllable.

We next applied classification of 96 hrs DE morphotype using the collected parameter data. We learned decision trees of depths 2 and 3, for predicting discrete morphotypes (GB, GT, GBT and DG). In order to assess feature importance and to avoid overfitting, we used a bootstrapping cross-validation approach, learning 500 models, each learned from randomly sampled 70% of the gastruloids in the experiment, and evaluated on the remaining 30% as a test set. We then examined the tree topologies that repeated most frequently, in classifiers that obtained test set accuracy above a set threshold (Fig. 2a, Fig. S2a). Examining the top decision node in these trees, we find that onset time of Sox17 was the most frequent top parameter, followed by gastruloid area (or size) at 72 hrs (Fig. 2b). The sample 4-class morphotype decision tree shown in Fig. 2c represents the most frequent tree topology. This tree shows that late Sox17 onset mostly leads to dispersed gut (DG), where a subset of gastruloids with more circular morphology at 72 hrs generate a gut tube (GT). For gastruloids with an early enough Sox17 onset time, smaller size at 72 hrs is predictive of GBT, while larger size gastruloids mostly end up with a GB morphotype. Thus, at some combinations of parameter ranges, morphotype outcome can be predicted at higher confidence, while in other ranges prediction is less definitive.

**Figure 2.**
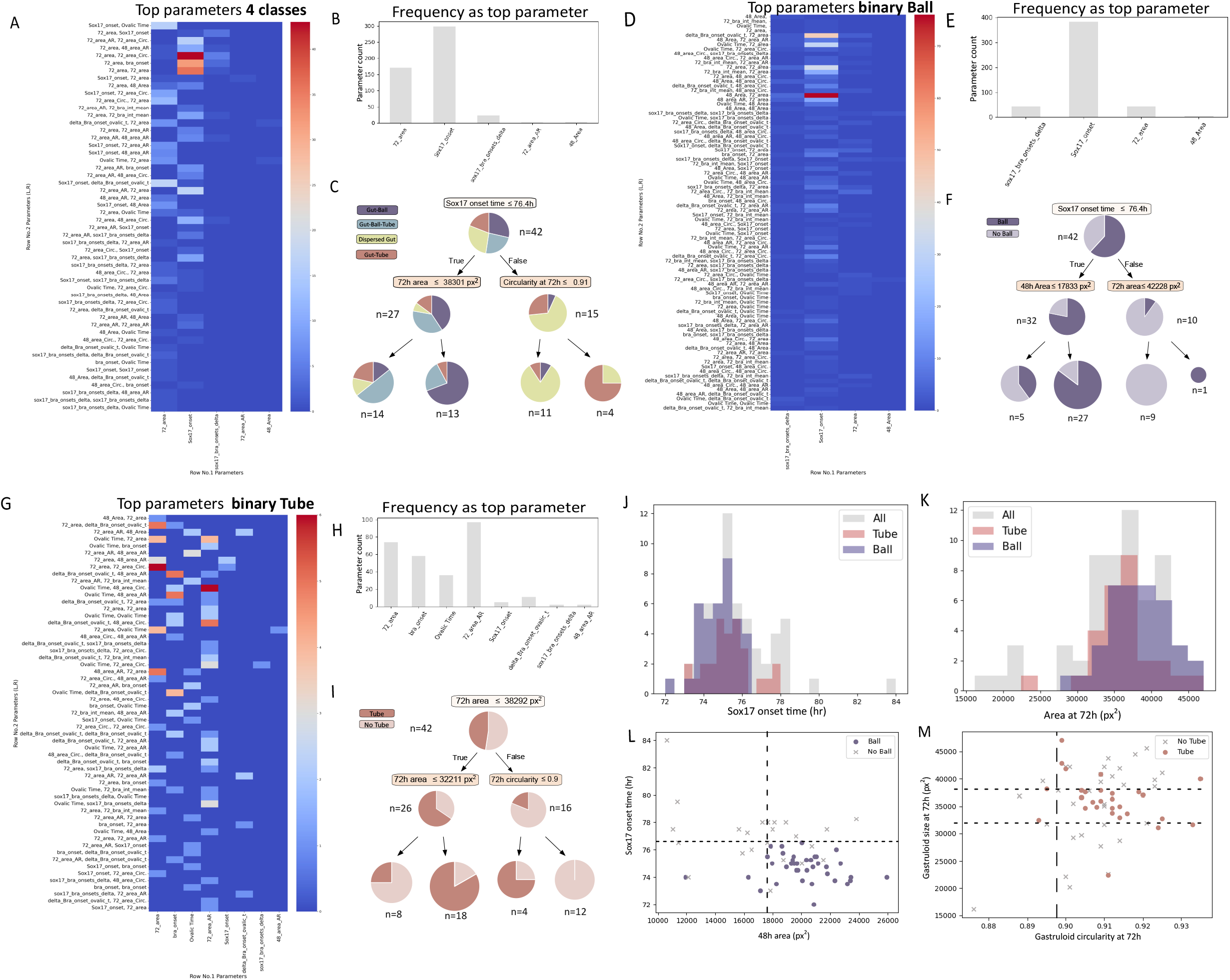
Early gastruloid morphology predicts endoderm morpho-type. **A**. Frequencies of decision tree topologies, within 500 trees learned in 4-morphotype classification (Gut-Ball, Gut-Tube, Gut-Ball-Tube and Dispersed-Gut). Each of the 500 trees was learned over 70% of the data, and tested over the remaining 30% (n=60). **B**. Frequency of top node parameters over the 500 learned trees. **C**. Sample tree for 4-class classification. **D**. Frequencies of decision tree topologies, learned in a binary Ball appearance classification task (n=60). **E**. Frequency of top node parameters for binary Ball classification. **F**. Sample tree for binary Ball classification. **G**. Frequencies of decision tree topologies, learned in a binary Tube appearance classification task (n=60). **H**. Frequency of top node parameters for binary Tube classification. **I**. Sample tree for binary Tube classification. **J**. Sox17 onset time distribution of Tube and Ball containing morphologies. **K**. 72h area distribution of Tube and Ball containing morphologies. **L-M**. Individual gastruloids plotted according to first and second level parameters of the sample Ball (L) and Tube(M) predicting trees.

Since the ball and tube morphologies can co-occur in the same gastruloid (in GBT gastruloids), we hypothesized that learning binary classifier trees, predicting any formation of Ball or Tube, may achieve better predictive power. The reasoning is that by binarizing the decision, we may direct the trees to focus on different key parameters that are essential for the formation of either ball or tube, independently. Fig. 2f,i show sample trees from each classifier type, representing the most abundant tree topology in each.

For binary Ball prediction, tree topologies with Sox17 onset time as a top parameter dominate, where second tier parameters only slightly refine the prediction (Fig. 2d,e,f). These predictors obtained higher test-set accuracy scores than the 4-class or binary Tube predictors (Fig. S2a). The most frequent tree topology suggests that early Sox17 onset, in combination with large size at 48 hrs will most likely lead to ball formation (Fig. 2j,l).

For binary Tube prediction, only few of the 2-level trees pass the accuracy threshold, and no single topology (or top level parameter) dominate the result (Fig. 2g,h). The sample tree that predicts for tube appearance suggests that a certain size range at 72 hrs leads to tube formation, while larger gastruloids will only generate a tube if they already started to lose their spherical shape at 72 hrs (Fig. 2I,k,m). Other common topologies either rely solely on ovoid-shape related parameters, or on combinations of size and Sox17 levels (Fig. S2d). Three-level trees did not obtain higher accuracy, nor did they focus on fewer top parameters (Fig. S2b,e). In summary, the poorer performance of Tube prediction suggests that the tube morphology is more complicated to predict, and its formation may depend on multiple factors. In contrast, binary prediction of Ball appearance showed better performance, with only a few frequent tree topologies, suggesting a simpler prediction task.

### Temporary delay in endoderm progression increases tube frequency

We next set to understand the dependency of the GB morphotype on early Sox17 onset. Gastruloids start elongating towards the end of Day 4. As Sox17+ cells sort out of the Bra+ population, and then upregulate E-cad to create epithelial domains (24), we hypothesized that in GB morphology this epithelialization occurs before the elongation process, thus leading to formation of tight epithelia that fail to migrate later along the elongating axis. Thus, the stage of endoderm progression at the start of gastruloid elongation may be critical to the resulting morphotype. Too advanced E-Cad+ endodermal cells may lead to Ball morphotype, while less advanced endoderm cells may display higher motility during elongation, allowing the formation of an organized tube (Movie S4).

To test this hypothesis, we generated a short delay in endoderm progression. As the Nodal/Activin pathway promotes the differentiation of the DE from the mesendoderm layer (24–27), we hypothesized that a temporary inhibition of this pathway could cause a short delay in the progression of endodermal differentiation, so that the majority of DE cells do not undergo MET until after the gastruloid substantially elongates, thus shifting DE morphology towards other narrow/tube containing patterns. Using the small molecule SB-431542 (‘SB’), a nodal-pathway inhibitor via activin receptor, we partially inhibited endoderm onset and/or expansion. We sought to find the optimal inhibition window that will generate the desired delay while still maintaining sufficient endodermal cells to generate a robust tube morphology.

When replacing CHIR with SB exposure at 66h (6 hours after Bra onset), both early long and short exposures to SB (12 and 6 hr, respectively) showed similarly low intensity of Sox17-RFP(+) cells compared to control (Fig. 3a,b). This early inhibition has a long lasting effect on the quantity of Sox17-RFP(+) cells, even after the inhibition pulse is over, and there is no recovery of the DE layer (Fig. 3c,d, S3a). These Nodal inhibition pulses have the opposite effect on mesoderm progression, marked by Bra-GFP intensity. The earlier SB pulse results in a higher Bra intensity then the later pulse and control (Fig. 3e). Similar to the effect on Sox17 expression, this effect is also long-lasting (Fig. 3f,g, S3b). This suggests that following inhibition of Nodal, Bra(+) mesendodermal progenitors continue to differentiate to mesodermal fates, and lost their potential to become DE. In contrast to the effect of early pulses of SB, a later, short pulse of SB (starting after the 24h CHIR pulse and 12h after Bra onset time) did not harm Sox17(+) cells onset and expansion (Fig.3b). However, this inhibition timing resulted in significant reduction of DE-domain diameter compared to the control at 96h (Fig. 3h). Under this protocol, morphotype distribution displays an increase in Tube appearance (75% compared to 63%, Fig. 3a). Notably, gastruloid ability to elongate is not affected by SB pulses (Fig. S3d). In summary, a late short inhibition of Nodal/Activin has the potential to increase the frequency of Tube morphotypes and reduce the endoderm domain diameter in all morphotypes, while not interfering with overall gastruloid morphology.

**Figure 3.**
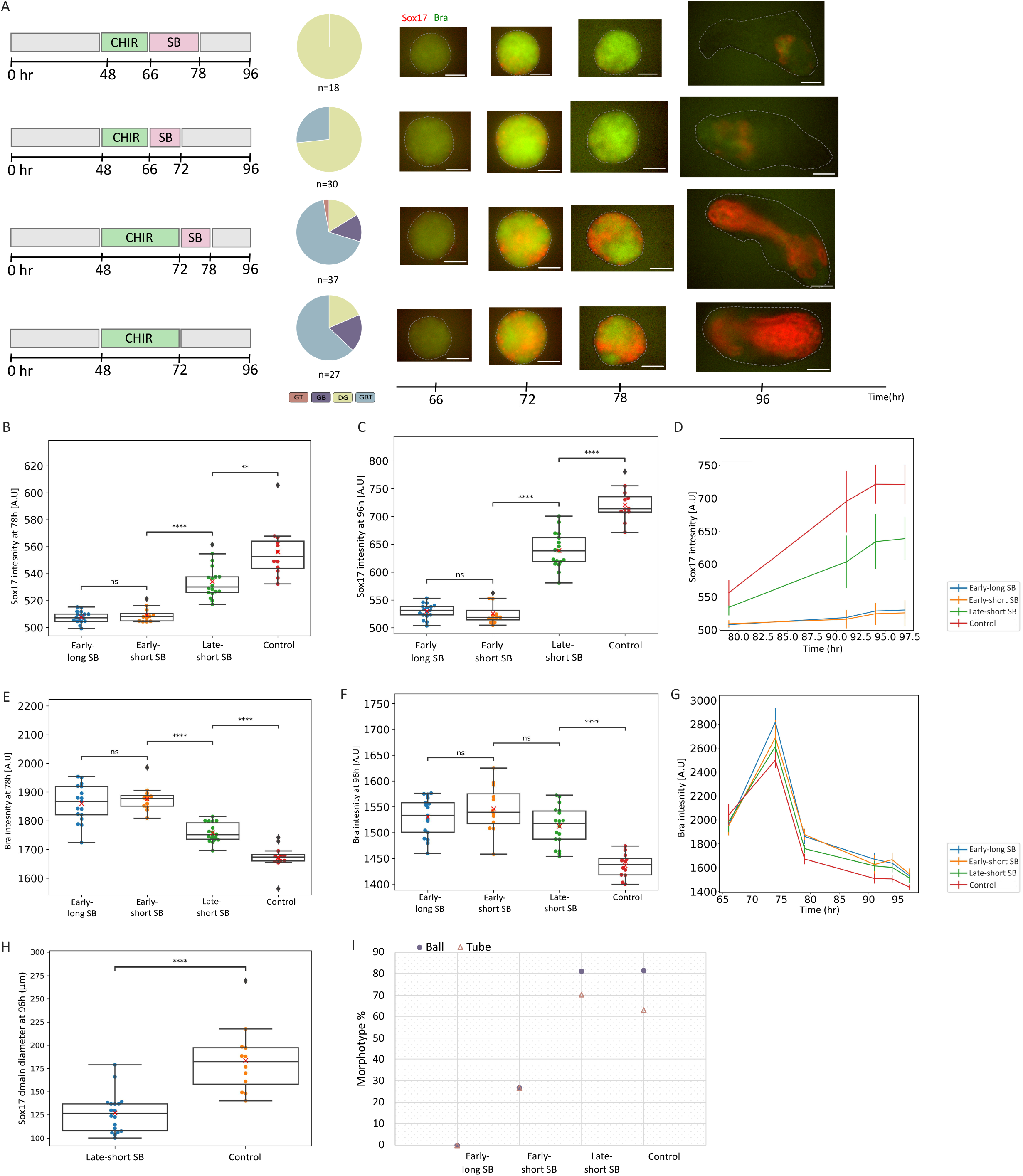
Short Nodal pathway inhibition pulse increases Tube frequency. **A**. SB treatment timelines: early-short, early-long or late-short SB treatments. Morphotypes distribution and sample time-line images (66,72,78,96h) for every condition. **B-C**. Average gastruloid Sox17-RFP levels at 78h (b) and 96h (c) under the different conditions. **D**. Sox17-RFP levels over time in each condition. **E-F**. Average gastruloid Bra-GFP levels at 78h (e) and 96h (f) under the different conditions. **G**. Bra-GFP levels over time in each condition. **F**. Sox17-RFP domain diameter at 96h for late-short SB and control groups. **I**. Frequency of Ball or Tube containing morphotypes at 96h for each condition.

### Timing gastruloid-specific external cue by gastruloid parameters reduces morpho-type variability

We have previously demonstrated that embryoid bodies (EBs) differ from each other in their differentiation rate, within and between experiments (19). This variability is also seen in the gastruloid basic differentiation protocol, for example when comparing Brachyury onset time (Fig. S2c). Since this protocol, like most others, relies on introduction of external signals at absolute times (e.g. addition of CHIR at 48hr), it is likely that each gastruloid is in a slightly different state at the time of signal addition. The interaction between internal temporal variability and uniformly-timed external signals can result in increased divergence of phenotypic outcome, as demonstrated in microorganism populations, where such divergence may be leveraged as a bet-hedging strategy (28–30). However, in the gastruloid system this may lead to undesired variability. In particular, the addition of a low-percentage extra-cellular matrix (Matrigel) to the medium at 96 hrs drives an extended elongation in gastruloids, as well as the formation of additional developmental structures such as somites. As in all steps of the protocol, Matrigel (MG) addition is applied to all gastruloids at the same time, regardless of their morphology or inner differentiation state. We hypothesized that a more “personalized”, gastruloid-based, addition of MG can line up the gastruloids within a narrower morpho-developmental window, resulting in a more robust DE outcome. This is further supported by the classifier analysis, where gastruloid shape descriptors (AR, circularity) are frequently selected for Tube appearance predictors (Fig. 2g,h). We therefore chose the gastruloid Aspect-Ratio (AR, width/length of the gastruloid), representing the elongation state, as the internal parameter to which we match the external signal.

MG was added to selected groups of gastruloids based on their oval shape (AR >= 1.2) at specific time points. Three MG addition times were tested, 5 hours apart (Fig. 4a). In the last addition time point (95h), MG was added to all the gastruloids that were left, regardless of their shape. In the control plate, MG was added in the same three addition times, but gastruloids were assigned randomly to each addition group (i.e. independent of their shape). To account for the effect of matrigel, DE morphotypes were estimated at 100 hrs.

**Figure 4.**
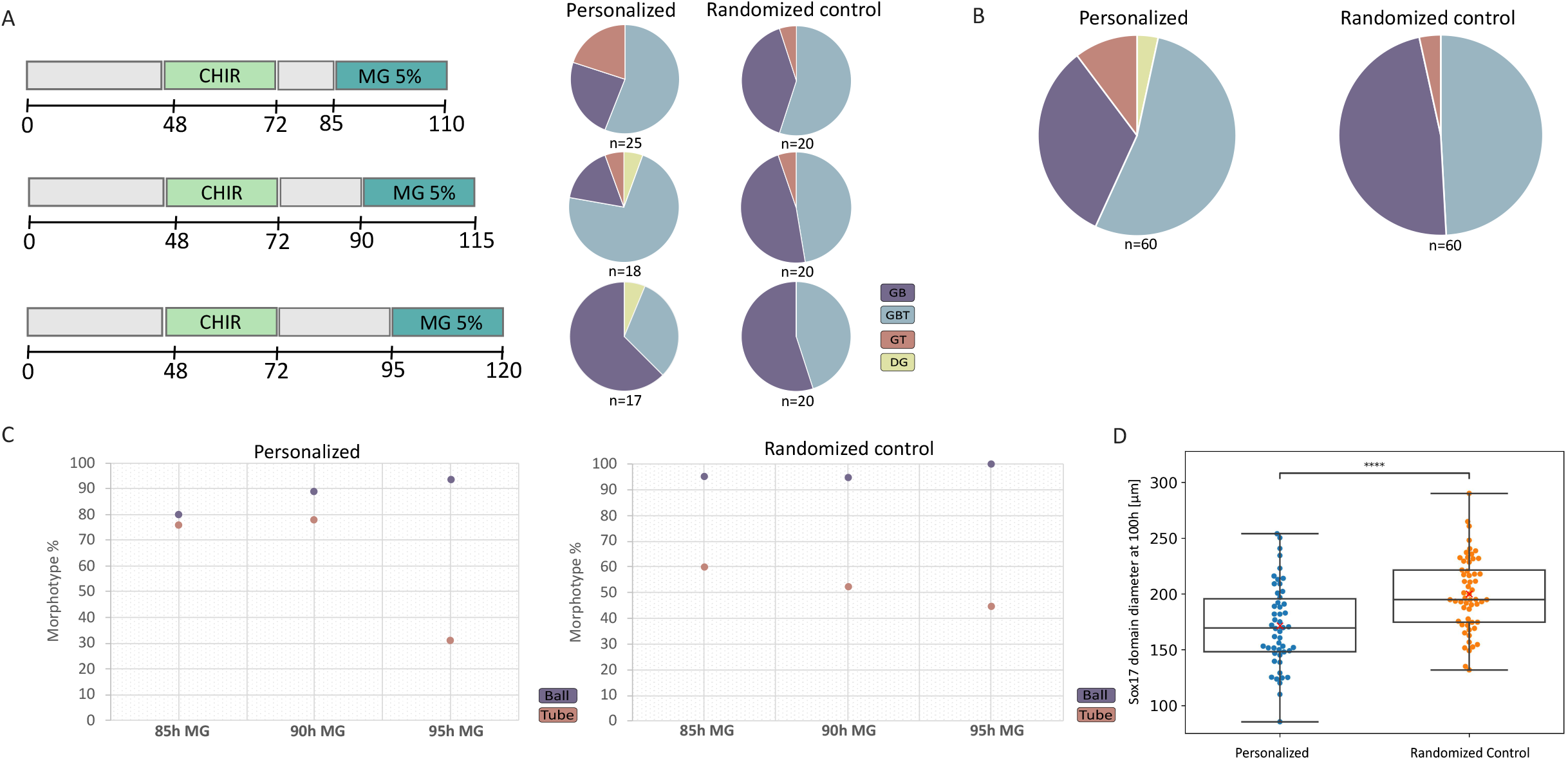
Gastruloid-specific timing of external cue by gastruloid parameters increases Tube frequency. **A**. Matrigel addition timelines and morphotype distribution for each addition time, for the personalized (gastruloid-specific) or randomized addition groups. **B**. Overall morphotype distribution for personalized and randomized MG addition plates. **C**. Binary Ball and Tube appearances in every addition time, in personalized (left) and randomized control (right) groups. **D**. Sox17 domain diameter for Ball containing gastruloids (GB and GBT), in the personalized and randomized control plates.

When comparing morphotype distribution for each MG addition-time group, we find that the two earlier addition groups (85h and 90h) obtained higher Tube rates compared to the last addition group (76%-78% vs. 31%), with a shift from GT to GBT in 90h (Fig. 4a,c). In contrast, GB rate dramatically increased in the last group (17%-24% vs. 62%). Notably, the control (randomized) gastruloids show, as expected, more similar distributions in all addition groups (Fig. 4a,c). In the first and second addition groups, personalized gastruloid selection increases Tube appearance to 76%-78% compared to 60%-52% in the matched groups of the randomized control (Fig. 4c). Moreover, regardless of morphotype distribution, personalized MG addition resulted in a smaller endoderm domain diameter (Fig. 4d).

These results indicate that personalization of MG addition could change the statistics of morphotype appearance. Combining such personalization with e.g. selection of early elongating gastruloids (groups 1 and 2 in Fig. 4a) can further enrich for a tube-containing morphotype.

## Discussion

In vitro models of early embryonic development using mouse embryonic stem cells are becoming increasingly popular as a tool to understand early developmental processes in the embryo. They allow looking at early processes outside of the uterus, manipulating them using external cues and following the dynamics of specific fluorescently-tagged marker genes. These models also have many limitations, a major one being the reproducibility and variability within and between experiments. Here we tackle the variability problem using endoderm development in gastruloids as a model. We focus on understanding the different endoderm morphologies in gastruloids, predicting each morphotype using early measurable parameters and trying to control its frequency, toward an overall increase of reproducibility and improved control in obtaining structures more similar to the embryo (in our case, meaning gut-tube like structures).

The high divergence of endoderm patterns and morphology within and between experiments can be explained by the combination of externally-timed cues in the protocol and different internal developmental time in each gastruloid. Developmental time can be influenced by aggregation time, initial aggregate size, initial pluripotency state of the cells, and more. Different developmental rates can also be seen between different cell lines, and even between different batches of media components. Given the ability to measure different descriptive parameters during live imaging of gastruloid development, we were able to predict certain morphologies based on specific parameters. Notably, even when the overall performance of the decision tree classifiers is not very high, we can draw biological insights from specific branches in those trees, that point at a specific morphotype and parameter range. Though we are naturally interested in parameter values that promote Tube appearance, understanding the parameters leading to other morphologies is also helpful, as we can intervene to change these parameters. Some of the predictive parameters, such as early gastruloid size which is highly predictive of Tube appearance, cannot be directly intervened with during the protocol. However, they may guide future modifications to the protocol, such as starting cell number, or adding a size-exclusion step. Other parameters, such as differentiation progression as measured by fluorescent markers, can suggest real-time interference, either uniformly to all gastruloids, or individually based on per-gastruloid measured parameters.

Our results point to two types of coordination required for gut tube structures to correctly emerge. The first type is between the gastruloid internal tempo and the external cues supplied in the medium. Embryo-like models have a natural variability in their rate of progression (19). Such temporal variability, combined with uniform external cues can lead to phenotypic variability between individuals (28,29,31). In particular, in the TLS/gastruloid protocol, there is some variability in the time the gastruloids start to obtain an ovoid shape. The “personalized” experiment demonstrated that matching the Matrigel addition time to the gastruloid ovoid-shape time results in increased tube frequency. The concept of matching external cue timing to gastruloid tempo may be applied to other phases of the protocol, such as CHIR addition or removal times, and also generalized to other organoid protocols.

The second coordination is between the endoderm progression rate and the gastruloid elongation process. Our results show that slightly delaying DE progression using a pulse of Nodal pathway inhibition, increases the frequency of tube-like structures. We have recently shown in both 2D and 3D cultures that DE cells switch on Sox17 while still in the mesenchymal state, and only fully turn up E-cadherin again after a period of expansion and self-sorting (24). In embryoid bodies this mesenchymal-to-epithelial transition leads to the formation of lumens or clusters. In the gastruloid protocol, multiple lumens join into the “gut-ball” mesh of lumens. If the gastruloid starts to elongate while DE cells are still mesenchymal, they can “hitchhike” on the elongation process and spread along the anterior-posterior axis of the gastruloid, before epithelializing into a tube (Movie S4). Here, the lack of inherent synchronization between these two processes may stem from the difference in DE morphogenesis between the gastruloid and the embryo. In the embryo, DE cells first intercalate into the surrounding PE layer, which in turn forms a tube in a folding process (6,32,33). In general, embryo-like models may display variability due to such lack of synchronization between two processes which in the embryo are coordinated through a third element (tissue or structure) which is missing in the model. Here we suggest additive cues in the protocol to try and balance this coordination. An important aspect is the timing of Nodal pathway inhibition, that should be finely tuned. In vitro mesoderm and endoderm fates are dependent on mesendoderm layer progression, and inhibition of Nodal disrupts and changes cell-fate decisions, where earlier inhibition timing pushed more cells toward mesodermal fates, up to complete ablation of the endoderm (Fig. 3). This suggests that it may be better to synchronize the inhibition timing with Bra onset time (in our case, around 12h after Bra onset), to obtain the right delay in endoderm progression with respect to mesodermal progression, which plays a more dominant role in driving elongation. In the future, this global inhibition can also be “personalized” based on gastruloid-specific Bra onset time, resulting in better control over endodermal morphologies.

Overall, this study suggests both personalized and global interventions to the gastruloid protocol that may direct endoderm to enrich for desired morphotypes. The approach we take here, based on analyzing morphotype predictive parameters, may be generalized to other tissue morphologies in the gastruloid system, as well as to other organoid protocols where morphotype variability poses an obstacle.

## Acknowledgements

We thank J. Veenvliet for helpful discussions, and members of our lab for comments on the manuscript. This study was supported by the Israel Science Foundation (1491/22), the European Union (Horizon-EIC-2021-PathfinderChallenges-01 101071203, SUMO) and the Good Food Institute research program.

## Author Contributions

NF and IN conceived the study. NF performed all experiments. NF and CS analyzed the data. NF and IN wrote the manuscript.

## Methods

### Cell culture

All experiments were conducted using a double-reporter mouse ESC line of Bra-GFP and Sox17-RFP {Pour 2022}. Cells were cultured on 0.2% gelatin coated plates on inactivated mouse embryonic fibroblast feeder cells at 37°C and 5% CO2 incubators. To maintain pluripotency, cells were cultured in ES-LIF medium containing knockout DMEM with 15% Fetal bovine serum (FBS) (BI, 04-001-1A), 1% penicillin/streptomycin (BI, 03-033-1B), 1% L-Glutamine (BI, 03-020-1B), 1% non-essential amino acids (BI, 01-340-1B), ß-mercaptoethanol and 1000 U/ml of leukemia inhibitory factor (LIF). For passaging, cells were detached from the plate by 3-minute incubation in 0.05% trypsin-EDTA (BI, 03-052-1A), followed by neutralization using equal amounts of growing medium (ES-LIF). Cells were then passed into a collection tube and centrifuged at 1000g for 4 minutes. The supernatant was then discarded and the cell’s pellet has re-suspended in ES-LIF media and seeded in a new well at a 1:10 ratio.

### Gastruloid differentiation assay

Gastruloid differentiation medium was based on the N2B27 medium, containing 50% DMEM/F-12 (Thermo Fisher, 21041-025) and 50% Neurobasal A liquid (Thermo Fisher, 21103-049), supplemented with 2 mM L-glutamine, 0.1 mM ß-mercaptoethanol, 0.033% BSA (Sigma, A8412), 0.5% N2 supplement (Thermo Fisher, 17502-048) and 1% B27 supplement (Thermo Fisher, 17504-044). For the creation of aggregates, cells were detached from the wells with trypsin as described above. After centrifugation, the cells were counted and diluted in N2B27 medium for the correct number of cells per ml needed for the creation of aggregates of a specific initial size (for 100-cell aggregates, the dilution was to 2500 cells/ml). Then a 40μl volume of the diluted cells were seeded in low-adherence 96 U-bottom wells (Greiner Bio-One, 650970) for 48h. At 48h, each well was supplemented with 110ul of N2B27+ Chiron (CHIR99021, BioVision, 1677-5), to a final CHIR concentration of 3μM. At a specified time, 5% Matrigel (Corning, 356231) was added to the cells, diluted in N2B27 (1ml media + 50μl matrigel 100%). For inhibition of the Nodal pathway, SB-431542 (Sigma *S4317)*, an Activin receptor inhibitor, was added at a final concentration of 10μM in N2B27 at indicated times. Aggregates were grown at all times in 37°C and 5% CO2 conditions.

### Live imaging

Images were taken using a Nikon TiE epi-fluorescence microscope equipped with a motorized XY stage (Prior) and an LED light source (Lumencor Sola), and taken with 20X or 10X magnification in up to three fluorescent wavelengths and phase contrast using NIS Elements software. The microscope is enclosed with an environmental chamber, maintaining 37°C and 5% CO2 (Okolab).

### Immunohistochemistry

For immunostainings, gastruloids were collected and washed with PBS x1 (Sigma D1408) with NaCl 8gr/L (Bio-Lab 19030591) and MgCl2 0.1gr/L (sigma 771-18-6) three times, and with PBS x1 one time. Gastruloids were transferred to PFA 4% and incubated on a shaking platform in 4°C for 1hr for fixation. Afterwards samples were washed with PBS x1 on a shaking platform twice, 5 min each time. Samples were then permeabilized with PBST (PBS, 0.5% Triton-x100 (sigma X-100), in room-temperature for 20min X 3 times. Samples were then incubated O.N. on a rocking platform in 4°C in Blocking Buffer (PBST, 5% FBS). Samples were then transferred to Primary ABs solution for 72h on rocking platform in 4°C. Primary ABs were diluted as recommended by manufacturer in blocking buffer. Samples were then washed X3 with blocking buffer and X3 with PBST, and were incubated in blocking buffer O.N. Samples were incubated with secondary ABs solution (diluted 1:1000 in blocking buffer) for 24hr. samples were then washed again with blocking buffer X3 and PBST X3 and kept in PBS 4°C until imaging. Antibodies used: E-cadherin (Cell Signaling 24E10), Alexa405 (secondary).

### Image analysis

To determine area (Size), lengths, and RFP or GFP fluorescent intensity of the samples, measurements were made using the Fiji program. The whole gastruloid contour and the endoderm region were traced manually. Parameter descriptions detailed in attached Table S1 and in Fig. S1

**Table S1:**

| Parameter | Measurment description | Units |
| --- | --- | --- |
| **Area** | Size of selected area. | Pixel |
| **Circularity** | Roundness level of selected area, defined as $4\pi \cdot \text{area} / \text{perimeter}^2$ .<br>Ranges from 0 to 1 (perfect circle) | |
| **Aspect-Ratio** | Major Axis/Minor Axis (L/D) of selected area. Ranges from 1 (perfect circles) - infinite. |  |
| **Intensity** | "Mean gray value" measurement. The average fluorescence intensity in the relevant channel within the selection area | A.U. |
| **Lengths** | using "straight line" selection tool in Fiji. | $\mu\text{m}$ |
| **Sox17 onset** | The earliest time point in which 15 Sox17-RFP+ cells are detected in the gastruloid | hours |
| **Bra onset** | The earliest time point in which mean gastruloid Bra-GFP intensity crossed a set threshold which depends on imaging parameters. | hours |

### Decision tree learning

Decision tree classifiers were learned from collected data using the python Scikit Learn package. Gini scores values represent a measurement of the impurity of the decision – the minimal gini score (0) means all cases fall in one category only. For four-class decision trees, the threshold test-set accuracy score for tree consideration was 0.4, while for binary classifier decision trees the threshold was set to 0.6.

**Figure S1.**
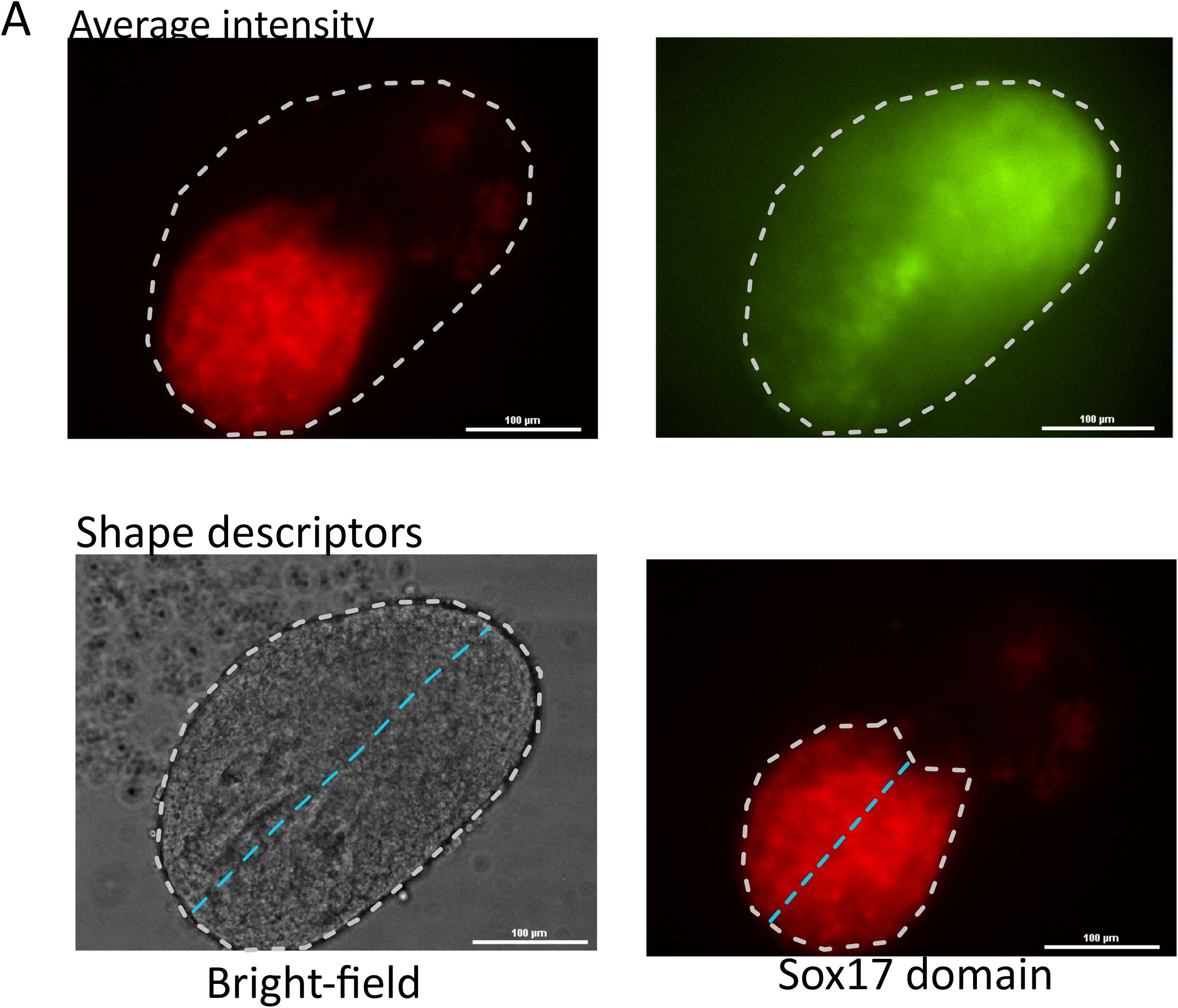
Morphometric parameter measurements. **A**. Selection area for whole gastruloid average intensity measurement. **B**. Shape descriptors measurements. Whole gastruloid (left, brigh-field) or Sox17-RFP domain (right, RFP channel) shape is shown by dashed while line (left). The shape was used for Area, AR, Circularity measurements. Lengths measurements example shown in light-blue dashed line.

**Figure S2.**
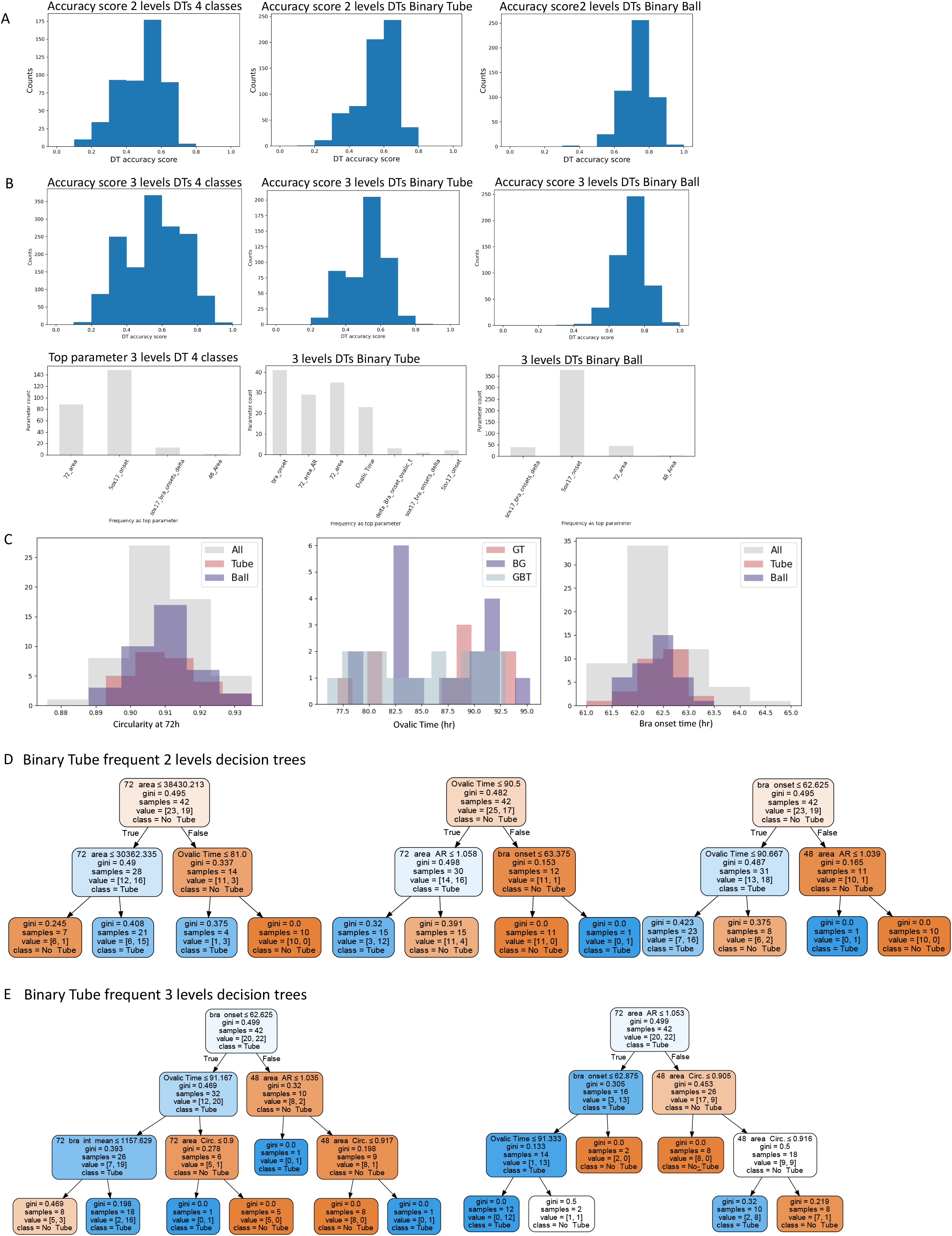
Decision tree statistics and sample trees. **A**. Test-set accuracy scores of 2-levels decision trees described in Fig. 2, for 4-classes, Tube and Ball classifications. **B**. Accuracy scores and Top frequent parameters of 3-levels decision trees, for 4-classes, Tube and Ball classifications. **C**. Left: Histograms of 72h circularity for Ball and Tube containing gastruloids, compared to all gastruloids. Right: Historgram of ovalic time (right) for GB,GBT and GT. **D**. Binary tube 2-levels decision tree examples for frequent top parameters. **E**. Binary tube 3-levels decision tree examples for frequent top parameters.

**Figure S3.**
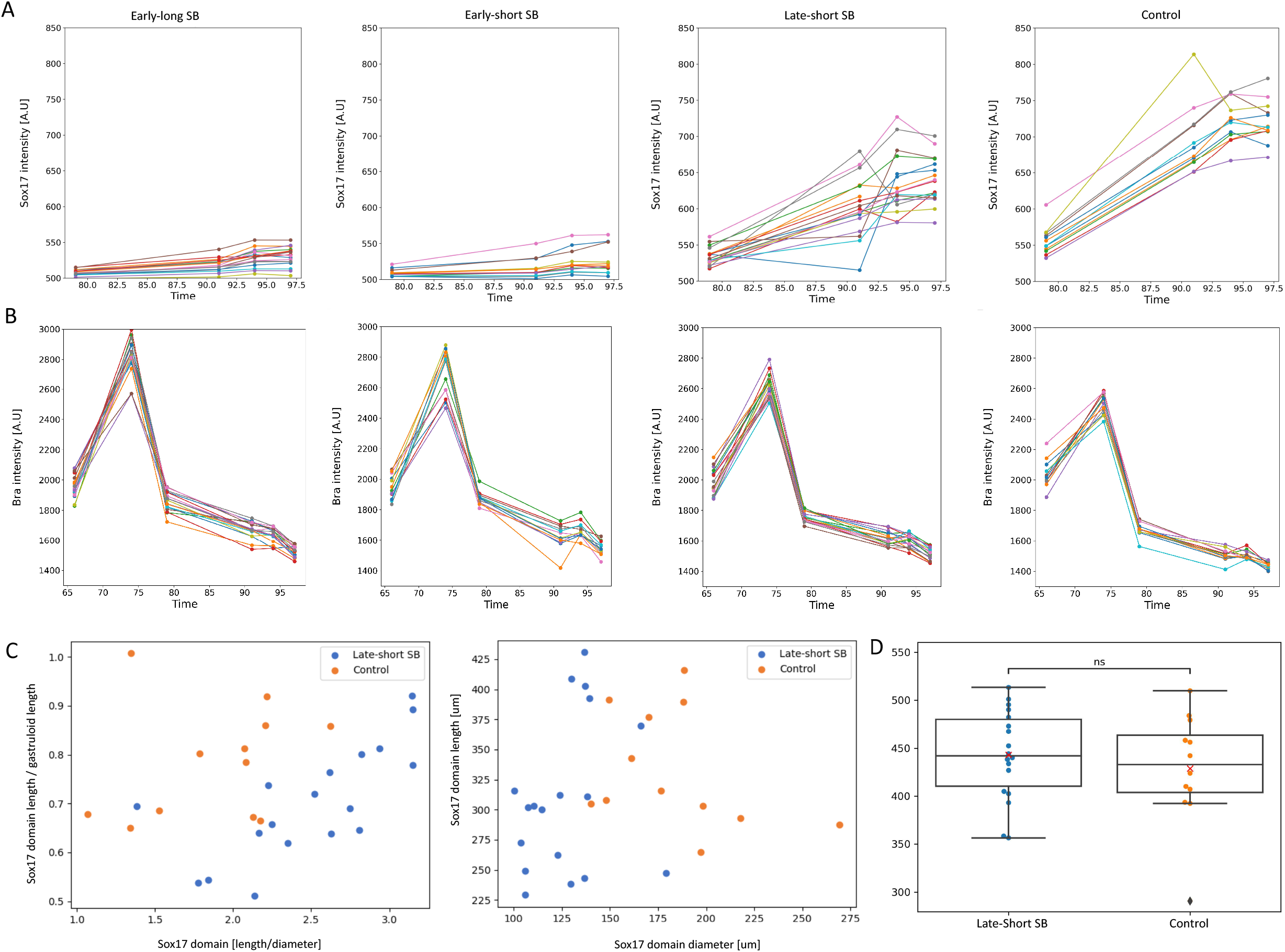
Nodal inhibition pulse effects on gastruloids. **A**. Sox17-RFP intensity over time for all SB conditions (left to right): early-long SB, early-short SB, late-short SB, control. Each track represents an individual gastruloid. **B**. Bra-GFP intensity over time for all SB conditions (left to right): early-long SB, early-short SB, late-short SB, control. **C**. Left: Comparison of late-short SB and control gastruloids according to dimensionless morphological parameters measured at 100h. Right: Comparison of Sox17-RFP domain length and diameter between Late-short SB and Control gastruloids.

**Movie S1**. Z-stack of a GB gastruloid at 96hrs, showing RFP channel (red) and DAPI (green). Interval between slices is 3um.

**Movie S2**. Time-lapse imaging of a GT gastruloid imaged between 100hrs and 120hrs, showing Sox17-RFP (red) and Bra-GFP (green).

**Movie S3**. Time-lapse imaging of a GBT gastruloid, imaged between 100hrs and 120hrs, showing Sox17-RFP (red).

**Movie S4**. Time-lapse imaging of an elongating GT gastruloid, imaged between 100hrs and 120hrs, showing the RFP channel.

## References

1. Tam PP. The control of somitogenesis in mouse embryos. J Embryol Exp Morphol. 1981 Oct;65 Suppl:103–28.

2. Jelier R, Kruger A, Swoger J, Zimmermann T, Lehner B. Compensatory Cell Movements Confer Robustness to Mechanical Deformation during Embryonic Development. Cell Syst. 2016 Aug 11;3(2):160–71.

3. Saiz N, Mora-Bitria L, Rahman S, George H, Herder JP, Garcia-Ojalvo J, et al. Growth-factor-mediated coupling between lineage size and cell fate choice underlies robustness of mammalian development. Elife. 2020 Jul 28;9.

4. Chal J, Pourquié O. Making muscle: skeletal myogenesis in vivo and in vitro. Development. 2017 Jun 15;144(12):2104–22.

5. Hubaud A, Pourquié O. Signalling dynamics in vertebrate segmentation. Nat Rev Mol Cell Biol. 2014 Nov;15(11):709–21.

6. Kwon GS, Viotti M, Hadjantonakis A-K. The endoderm of the mouse embryo arises by dynamic widespread intercalation of embryonic and extraembryonic lineages. Dev Cell. 2008 Oct;15(4):509–20.

7. Nowotschin S, Setty M, Kuo Y-Y, Liu V, Garg V, Sharma R, et al. The emergent landscape of the mouse gut endoderm at single-cell resolution. Nature. 2019 May;569(7756):361–7.

8. Oates AC, Morelli LG, Ares S. Patterning embryos with oscillations: structure, function and dynamics of the vertebrate segmentation clock. Development. 2012 Feb;139(4):625–39.

9. Saiz N, Hadjantonakis A-K. Coordination between patterning and morphogenesis ensures robustness during mouse development. Philos Trans R Soc Lond B, Biol Sci. 2020 Oct 12;375(1809):20190562.

10. Viotti M, Nowotschin S, Hadjantonakis A-K. SOX17 links gut endoderm morphogenesis and germ layer segregation. Nat Cell Biol. 2014 Dec;16(12):1146–56.

11. van den Brink SC, Baillie-Johnson P, Balayo T, Hadjantonakis A-K, Nowotschin S, Turner DA, et al. Symmetry breaking, germ layer specification and axial organisation in aggregates of mouse embryonic stem cells. Development. 2014 Nov;141(22):4231–42.

12. Poh Y-C, Chen J, Hong Y, Yi H, Zhang S, Chen J, et al. Generation of organized germ layers from a single mouse embryonic stem cell. Nat Commun. 2014 May 30;5:4000.

13. Turner DA, Girgin M, Alonso-Crisostomo L, Trivedi V, Baillie-Johnson P, Glodowski CR, et al. Anteroposterior polarity and elongation in the absence of extra-embryonic tissues and of spatially localised signalling in gastruloids: mammalian embryonic organoids. Development. 2017 Nov 1;144(21):3894–906.

14. Warmflash A, Sorre B, Etoc F, Siggia ED, Brivanlou AH. A method to recapitulate early embryonic spatial patterning in human embryonic stem cells. Nat Methods. 2014 Aug;11(8):847–54.

15. Beccari L, Moris N, Girgin M, Turner DA, Baillie-Johnson P, Cossy A-C, et al. Multi-axial self-organization properties of mouse embryonic stem cells into gastruloids. Nature. 2018 Oct 3;562(7726):272–6.

16. ten Berge D, Brugmann SA, Helms JA, Nusse R. Wnt and FGF signals interact to coordinate growth with cell fate specification during limb development. Development. 2008 Oct 1;135(19):3247–57.

17. van den Brink SC, Alemany A, van Batenburg V, Moris N, Blotenburg M, Vivié J, et al. Single-cell and spatial transcriptomics reveal somitogenesis in gastruloids. Nature. 2020 Jun;582(7812):405–9.

18. Veenvliet JV, Bolondi A, Kretzmer H, Haut L, Scholze-Wittler M, Schifferl D, et al. Mouse embryonic stem cells self-organize into trunk-like structures with neural tube and somites. BioRxiv. 2020 Mar 4;

19. Boxman J, Sagy N, Achanta S, Vadigepalli R, Nachman I. Integrated live imaging and molecular profiling of embryoid bodies reveals a synchronized progression of early differentiation. Sci Rep. 2016 Aug 17;6:31623.

20. Phipson B, Er PX, Combes AN, Forbes TA, Howden SE, Zappia L, et al. Evaluation of variability in human kidney organoids. Nat Methods. 2019 Jan;16(1):79–87.

21. Quadrato G, Nguyen T, Macosko EZ, Sherwood JL, Min Yang S, Berger DR, et al. Cell diversity and network dynamics in photosensitive human brain organoids. Nature. 2017 May 4;545(7652):48–53.

22. Vianello SD, Lutolf M. In vitro endoderm emergence and self-organisation in the absence of extraembryonic tissues and embryonic architecture. BioRxiv. 2020 Jun 9;

23. Anlas K, Trivedi V. Studying evolution of the primary body axis in vivo and in vitro. Elife. 2021 Aug 31;10.

24. Pour M, Kumar AS, Farag N, Bolondi A, Kretzmer H, Walther M, et al. Emergence and patterning dynamics of mouse-definitive endoderm. iScience. 2022 Jan 21;25(1):103556.

25. Camus A, Perea-Gomez A, Moreau A, Collignon J. Absence of Nodal signaling promotes precocious neural differentiation in the mouse embryo. Dev Biol. 2006 Jul 15;295(2):743– 55.

26. Kubo A, Shinozaki K, Shannon JM, Kouskoff V, Kennedy M, Woo S, et al. Development of definitive endoderm from embryonic stem cells in culture. Development. 2004 Apr;131(7):1651–62.

27. Zhou X, Sasaki H, Lowe L, Hogan BL, Kuehn MR. Nodal is a novel TGF-beta-like gene expressed in the mouse node during gastrulation. Nature. 1993 Feb 11;361(6412):543–7.

28. Nachman I, Regev A, Ramanathan S. Dissecting timing variability in yeast meiosis. Cell. 2007 Nov 2;131(3):544–56.

29. Yurkovsky E, Nachman I. Event timing at the single-cell level. Brief Funct Genomics. 2013 Mar;12(2):90–8.

30. Fujita M, Losick R. Evidence that entry into sporulation in Bacillus subtilis is governed by a gradual increase in the level and activity of the master regulator Spo0A. Genes Dev. 2005 Sep 15;19(18):2236–44.

31. Doncic A, Falleur-Fettig M, Skotheim JM. Distinct interactions select and maintain a specific cell fate. Mol Cell. 2011 Aug 19;43(4):528–39.

32. Burtscher I, Lickert H. Foxa2 regulates polarity and epithelialization in the endoderm germ layer of the mouse embryo. Development. 2009 Mar;136(6):1029–38.

33. Rivera-Pérez JA, Hadjantonakis A-K. The dynamics of morphogenesis in the early mouse embryo. Cold Spring Harb Perspect Biol. 2014 Jun 26;7(11).

